# The *SMYD1* p.Asn101Ser is a partial loss-of-function variant that impairs mitochondrial function and leads to early-onset cardiomyopathy

**DOI:** 10.64898/2026.07.28.741372

**Authors:** Marta W. Szulik, Clint Gwynn, Magnus Creed, Lina Ghaloul-Gonzalez, Sarah Franklin

## Abstract

Infantile cardiomyopathies are rare, life-threatening disorders for which genetic diagnosis has been accelerated by next-generation sequencing approaches, including gene panel, exome, and genome sequencing. However, determining the functional consequences of identified variants remains a major challenge. Variants in *SMYD1*, a striated muscle-specific lysine methyltransferase critical for cardiac development and mitochondrial function, have only recently been linked to human cardiomyopathy. Here, we functionally characterize a homozygous *SMYD1* variant (c.302A>G; p.Asn101Ser) identified in a patient with severe early-onset cardiomyopathy requiring cardiac transplantation. Structural modeling predicts that the N101S substitution perturbs a highly conserved residue near the cofactor binding pocket within SMYD1’s catalytic domain, disrupting local interactions and modestly destabilizing the protein. Consistent with these predictions, in vitro studies demonstrate that the N101S variant impairs mitochondrial respiratory capacity in myocytes. Quantification of SMYD1 protein levels in patient cardiac tissue revealed increased SMYD1 abundance, suggesting that the N101S variant results in functional impairment rather than protein instability and may trigger compensatory upregulation of SMYD1 expression. Together, these findings support a hypomorphic mechanism in which the N101S variant disrupts SMYD1 activity, leading to mitochondrial dysfunction and cardiomyopathy. This study provides mechanistic insight into SMYD1-associated cardiomyopathy and highlights the importance of integrating genetic, structural, and functional analyses to establish the pathogenicity of rare variants.

**New & Noteworthy:** This study provides the first functional characterization of the cardiomyopathy-associated SMYD1 N101S variant identified in a child with severe infantile cardiomyopathy. Structural modeling predicts reduced protein stability, while cellular assays demonstrate impaired mitochondrial respiratory function, supporting a hypomorphic effect. These findings establish a mechanistic link between SMYD1 dysfunction and infantile cardiomyopathy and highlight the importance of integrating genomic and functional approaches in rare cardiovascular disease.

## 1 Introduction

Infantile cardiomyopathies are rare but severe disorders associated with high morbidity and mortality, with a substantial proportion of affected children progressing to heart transplantation or death early in life (1, 2). Advances in genomic technologies, particularly next-generation sequencing approaches, including gene panels, exome, and genome sequencing (ES/GS), have significantly improved the identification of genetic variants underlying these conditions. However, establishing the functional consequences of identified variants remains a major challenge, particularly for rare or novel variants classified as variants of uncertain significance (VUS).

SMYD1 (SET and MYND domain-containing protein 1) is a lysine methyltransferase, which has been shown to methylate lysine 4 on histone H3, an established mark of gene activation (3–9). SMYD1 is expressed only in skeletal and cardiac muscle and has been shown to play a significant role in regulating cardiac development (10–12). Specifically, constitutive loss of *Smyd1* in mice *in utero* leads to death at embryonic stage E9.5 due to cardiac defects, manifested by disrupted maturation of ventricular cardiomyocytes, malformation of the right ventricle, and truncation of the outflow tract (10, 12). It has also been shown that conditional cardiomyocyte-specific loss of *Smyd1* is lethal at embryonic stage E12.5-15.5 accompanied by pericardial edema, thinned pericardium, and decreased trabeculation (12). In the adult myocardium, using inducible, cardiomyocyte-specific *Smyd1* knockout mice, loss of SMYD1 leads to massive downregulation of mitochondrial bioenergetics (9), cardiac hypertrophy, fibrosis, and heart failure (11). Despite extensive characterization of SMYD1 in zebrafish and mouse model systems, human data linking SMYD1 variants to disease and their molecular mechanisms remain limited. Only in the last few years have human patients been identified with variants in the *SMYD1* gene exhibiting cardiomyopathies. There are currently two previous reports of patients with *SMYD1* variant-associated cardiomyopathies: the first was a 24 years old patient with hypertrophic cardiomyopathy and a *de novo* heterozygous variant in the *SMYD1* gene (c.814T>C; p.Phe272Leu) (13); the second patient displayed left ventricular non-compaction (LVNC) cardiomyopathy and arrhythmias who underwent heart transplantation and has a truncating heterozygous variant in *SMYD1* gene (c.675delA; p.Lys225Asnfs*8) (14).

A third patient (P1) carrying a homozygous variant (c.302A>G; p.Asn101Ser) was previously described in a brief report in 2019 (15), and in a comprehensive clinical study in 2023 (16), which detailed the patient’s severe early-onset cardiomyopathy requiring mechanical circulatory support and cardiac transplantation. While these studies established the clinical phenotype and identified the variant through ES, the molecular and functional consequences of the N101S substitution remained unknown.

In the present study, we build upon these prior clinical observations and perform a functional characterization of the SMYD1 N101S variant. Using computational structural modeling, we demonstrate that this substitution is predicted to modestly destabilize the SMYD1 protein. Furthermore, using a site-directed mutagenesis approach and in vitro assays in cultured cells, we show that the N101S variant impairs mitochondrial respiratory function, consistent with a hypomorphic effect. Together, these findings provide mechanistic insight into the pathogenicity of the SMYD1 N101S variant and illustrate the importance of integrating genetic discovery with functional validation to better understand the molecular basis of cardiomyopathy.

## 2 Materials and methods

### 2.1 Patient and clinical data

Informed consent was obtained and approved for the patient in accordance with the University of Pittsburgh IRB-approved protocol #STUDY19040093. All methods were performed in accordance with the relevant guidelines and regulations outlined by the IRB.

A brief report of the proband P1 carrying the novel *SMYD1* variant was published in 2019 (15), and a comprehensive overview of the phenotype, including the proband’s original presentation, medical care requiring biventricular assist device (BiVAD) implant, and subsequent cardiac transplant, was published in 2023 (16). Notably, subsequent genetic analysis following heart transplantation revealed that P1 was also homozygous for a pathogenic *MYBPC3* variant (c.1224-52G>A; IVS13-52G>A), which is independently known to cause severe cardiomyopathy (16). Consequently, defining the individual and combined functional contributions of the *SMYD1* and *MYBPC3* variants will be essential for understanding disease pathogenesis.

The proband’s cardiac and skeletal muscle samples were acquired at the time of heart transplant, at which time the proband (P1) was three months of age. The age- and sex-matched control tissue samples were acquired from a 4-month-old female diagnosed with severe hypoxic encephalopathy and necrotizing enterocolitis (NEC).

### 2.2 Molecular modeling

We modeled the SMYD1 protein structure containing the N101S substitution using the published crystal structure of mouse SMYD1 in the PDB Database (PDB ID: 3n71) (17) in PyMOL (Schrodinger LLC). Our analysis of the SMYD1-N101S structure was performed using UCSF Chimera, developed by the Resource for Biocomputing, Visualization, and Information at the University of California, San Francisco (18). Molecular modeling simulation was performed by applying a series of structure minimization algorithms. Briefly, steepest descent minimization (100 steps, 0.02 Å step size) was performed first, followed by conjugate gradient minimization (100 steps, 0.02 Å step size) to reach an energy minimum. Minimization was performed *in vacuo*, with hydrogens and charges included, using Amber ff14SB force field (19).

To further evaluate the impact of the N101S substitution on protein stability and dynamics, we used the DynaMut2 web server (20). The crystal structure of SMYD1 (PDB ID: 3n71) was used as input, and the N101S substitution was introduced in silico. DynaMut2 integrates normal mode analysis with graph-based signatures to predict the effects of missense variants on protein stability and flexibility. The predicted change in Gibbs free energy (ΔΔG, kcal/mol) was used to assess the impact of the variant on structural stability, where negative values indicate destabilization and positive values indicate stabilization.

### 2.3 Site-directed mutagenesis

For all the cellular experiments carried out in this study, we utilized the mouse *Smyd1* isoform 1 (GenBank: NM_001160127.1), which is the mouse ortholog of human *SMYD1*. The full-length mouse *Smyd1* ORF cDNA tagged with MYC and FLAG on the C-terminus, cloned into the pCMV6-Entry vector, was purchased from OriGene. Site-directed mutagenesis was carried out using QuikChange II Site-Directed Mutagenesis kit (Agilent) according to the manufacturer’s instructions. To generate c.302A>G missense variant in the mouse *Smyd1* sequence, primers were designed using QuikChange Primer Design Tool (Agilent) (FWD: 5’-AATATGGGAAAGTGCCCAGCGAGAACATCAGGCTG-3’; REV: 5’-CAGCCTGATGTTCTCGCTGGGCACTTTCCCATATT-3’). To verify the presence of the N101S missense variant in the *Smyd1* sequence, plasmids containing the c.302A>G variant and unmodified plasmid controls were sequenced in the DNA sequencing core facility, University of Utah, and DNA Sanger sequencing data was analyzed using DNA Dynamo Sequence Analysis software.

### 2.4 Cell culture and transfection

Cultured C2C12 mouse myoblasts, undifferentiated, were plated in 60mm dishes at 60% confluency in growth medium (DMEM) supplemented with 10% FBS and 1% penicillin-streptomycin. Undifferentiated C2C12 cells do not express SMYD1; thus, they were transfected with either the *Smyd1*-N101S plasmid for expression of the N101S variant, SMYD1, or an empty vector as a transfection control (9). Lipofectamine 3000 and 7ug of plasmid were mixed in serum-free OPTI-MEM media and incubated at room temperature for 15 min. The plasmid DNA-lipofectamine complex was added to cell culture dishes. Non-transfected cells were used as an additional control. Cells were incubated for 48h before harvesting for endpoint experiments.

### 2.5 Cell Mito Stress Test

To evaluate mitochondrial function, the Cell Mito Stress Test was performed on undifferentiated C2C12 mouse myoblasts using a Seahorse XFe96 Flux Analyzer (Agilent), as previously described (9, 21). Briefly, the C2C12 cells were seeded in the 96-well Seahorse analyzer at a confluency of 20k cells/well. After ∼18h incubation, cells were transfected with *Smyd1*-N101S plasmid or wild-type *Smyd1* plasmid (as a control), using Lipofectamine 3000 in OPTI-MEM (Gibco) (9, 11). After 48h, measurements of the oxygen consumption rate (OCR) were conducted in XF base medium (Agilent-Seahorse) at pH 7.4 and supplemented with 2mM L-glutamine, 1mM pyruvate and 25mM glucose (n=6 replicates per condition). The OCR rates were measured before and after sequential addition of inhibitors: oligomycin (1µM), carbonyl cyanide 4-(trifluoromethoxy) phenylhydrazone (FCCP, 5µM), and mix of rotenone (1µM) and antimycin A (1µM). Following the OCR measurements, cells were stained with Hoechst 33342 to stain nuclei, and the nuclear signal intensity was measured using BioTek Cytation 5 (excitation 360nM, emission 450nm). The OCR values were normalized to the intensity of nuclear staining.

### 2.6 Electrophoresis and Western blotting

Protein separation and western blot analysis were carried out as previously described (9, 11, 21, 22). Briefly, protein samples were placed in Laemmli buffer, resolved on 10% SDS PAGE gels, and then transferred to nitrocellulose membranes using the semi-dry transfer method (Bio-Rad). The success of protein transfer to the membrane was confirmed using Ponceau S (0.1% w/v in 5% acetic acid). These membranes were blocked with 5% milk in TBS-T. Antibodies used in this study are as follows: Anti-SMYD1 (Abcam, ab32482), Anti-MYC (Abcam, ab9106), and Anti-PGC-1α (Abcam, an54481), followed by goat anti-rabbit secondary antibody (Abcam, ab97051). The signal was normalized to β-TUBULIN (Abcam, ab6046).

### 2.7 Gene expression analysis

Real-time qPCR analysis was performed as described previously (9, 11, 21, 22). Briefly, RNA was extracted using TRIzol (Invitrogen). Reverse transcription was performed from 1μg of mRNA using SuperScript First-Strand Synthesis system (Thermo Fisher Scientific). cDNA transcripts were amplified on a real-time PCR detection system (CFX Connect, Bio-Rad) with SYBR Green Supermix (Bio-Rad). The *Cq* values from the targeted genes were normalized by the gene expression level of *β-Actin* (FWD: TGTTACCAACTGGGACGA; REV: GGGGTGTTGAAGGTCTCA), and fold change was calculated using the ΔΔCt method. The following primers were used: *Smyd1* (FWD: CGAGGGTTTGTATCACGAGGTTGT; REV: CATGGGTCACCAGGAGAATAGCAT), *Ppargc1α* (FWD: CTTCGAGCTGTACTTTTGTGGACGGAA; REV: CTCTGAGCTTCCTTCAGTAAACTATCAAA), *Nppa* (FWD: CTGATGGATTTCAAGAACCTGCT; REV: CTCTGGGCTCCAATCCTGTC), *Myh6* (FWD: GAACAGCTGGGAGAAGGGGG; REV: GCCTCTGAGGCTATTCTATTGG).

### 2.8 Statistical Analysis

Unless specifically noted, data were collected from technical triplicates of at least three biological replicates of three independent experiments and presented as the means with standard error of the mean (SEM). Statistical significance was evaluated using the unpaired Student’s *t*-test or one-way ANOVA with multiple comparisons. An asterisk * that indicates *p*<0.05 was considered statistically significant. All biostatistics calculations were performed with Prism (GraphPad) or Excel (Microsoft Office).

## 3 Results

### 3.1 Summary of clinical presentation and genetic findings

The clinical presentation of the proband (P1) carrying the homozygous *SMYD1* c.302A>G (p.Asn101Ser) variant has been previously described in detail in our earlier reports (15, 16). Briefly, P1 was a female child of unaffected, healthy parents in a consanguineous relationship. The child was delivered at full term via scheduled C-section following an uncomplicated pregnancy to a mother who had previously experienced one miscarriage, one pregnancy that resulted in a stillborn child, one termination of pregnancy, and a healthy pregnancy with the same partner (Figure 1). The proband’s older male sibling was unaffected and presented with a normal echocardiogram, as did the parents. P1 presented with severe early-onset cardiomyopathy requiring biventricular assist device (BiVAD) support followed by cardiac transplantation in early infancy (15, 16).

**Figure 1.**
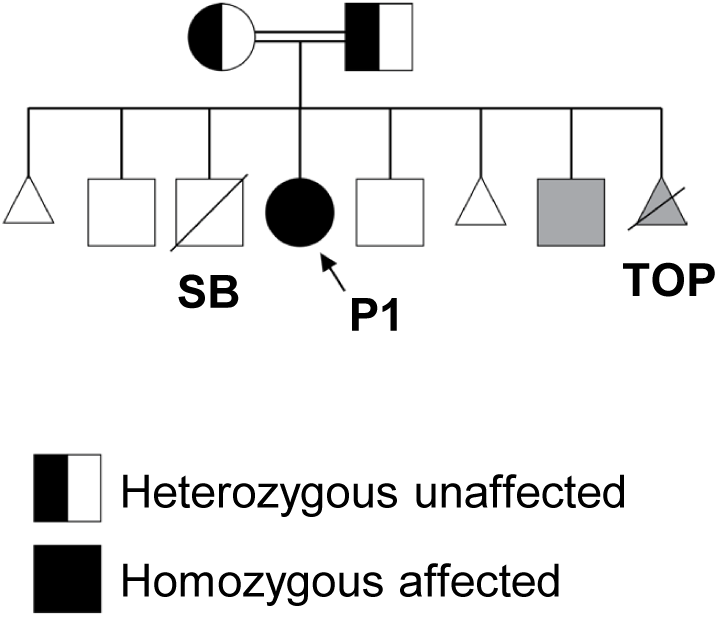
Pedigree of a family carrying a novel *SMYD1* variant (c.302A>G; p.Asn101Ser). Females are represented by circles and males by squares. Pregnancy losses are indicated as follows: miscarriages by triangles, stillbirth (SB) by a square with a diagonal line, termination of pregnancy (TOP) by a triangle with a diagonal line. A double line indicates consanguinity. Genotyped and negative for *SMYD1* variant are shown as grey symbols. Unaffected and ungenotyped individuals are shown as open symbols. The parents are unaffected and heterozygous for the variant, as indicated by half-filled symbols. The proband P1 (arrow) is homozygous and affected, indicated by a black circle.

Whole exome sequencing (WES) performed as part of the prior study identified the homozygous SMYD1 N101S variant, providing the basis for subsequent mechanistic investigation. For the purpose of the current study, cardiac and skeletal muscle tissues were obtained at the time of heart transplantation when P1 was three months of age. Age- and sex-matched control tissues were obtained from a four-month-old female donor.

### 3.2 Molecular modeling and prediction of protein structure perturbations

SMYD1 contains five functional domains based on sequence and structure homology to other methyltransferases, including: 1) the N-terminal SET catalytic domain (green), which is split by a 2) MYND (blue) domain and is followed by a 3) SET-I domain (green), 4) post-SET domain (green), and 5) C-terminal domain (CTD, orange) (Figure 2A). The split SET domain has two regions: the S sequence, implicated in cofactor binding, protein-protein interactions, and structural stability, and the ET domain, which contributes to SMYD1’s enzymatic activity (17). The SET domain is further supported by the “SET-I” and “post-SET” domains, both of which participate in cofactor and substrate binding. Previously reported *SMYD1* variants associated with cardiomyopathy localized to the ET and post-SET domains (13, 14).

**Figure 2.**
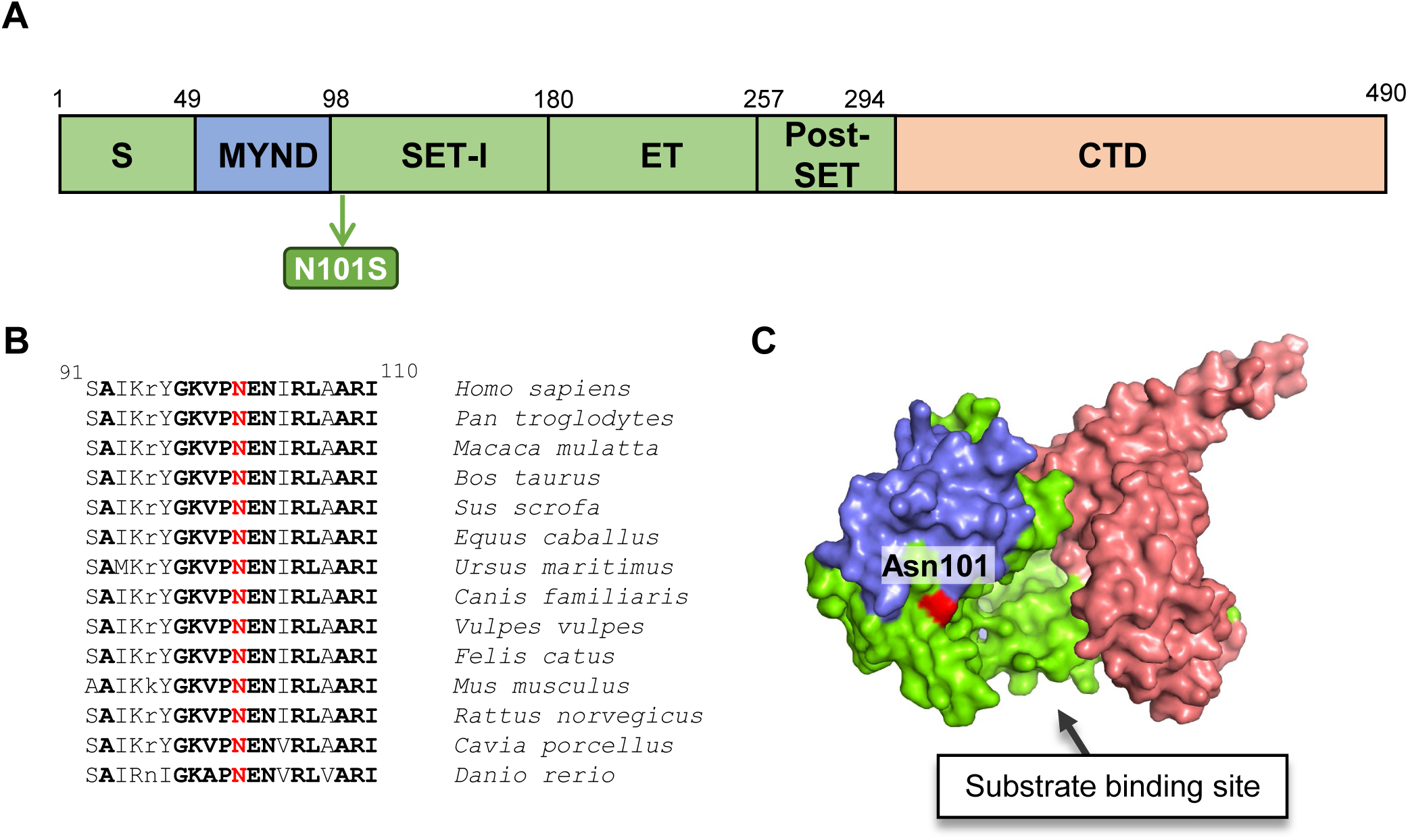
Structural domain organization of SMYD1 and identification of a novel *SMYD1* (c.302A>G; p.Asn101Ser) variant. **A.** Linear representation of the predicted structural domains of the SMYD1 protein in humans based on published mouse SMYD1 domains. A single amino acid substitution at position 101 (Asn to Ser) was identified in a pediatric patient (P1) with biventricular heart failure as indicated with a green arrow. **B.** SMYD1 protein sequence alignment from position 91 through position 110 spanning the MYND-SET-I domain boundary, showing the conserved Asn101 residue (highlighted in red) across 14 vertebrate species. Residues in bold capital letters are invariant across all 14 species, residues in capital letters (not bold) are highly conserved, and residues in lower case letters appear to be dispensable. **C.** Crystal structure of mouse SMYD1 (PDB ID: 3N71) showing the location of residue Asn101 (red) near the substrate binding site (black arrow).

The *SMYD1* variant (c.302A>G; p.Asn101Ser) is located within the SET-I domain at its N-terminus (Figure 2A). This single nucleotide substitution at position 302 from adenine to guanine (c.302A>G) results in an amino acid change from asparagine to serine at position 101 (abbreviated Asn101Ser or N101S). Sequence alignment demonstrates that Asn101 is highly conserved in SMYD1 across species (Figure 2B), supporting a potential functional role. In humans, SMYD1 protein in cardiac muscle shares 94% sequence homology with one of two muscle-specific transcripts in mice, making mouse SMYD1 isoform 1 the ortholog to human SMYD1 (23). Structural data from mouse SMYD1 indicate a deep, partially enclosed cofactor binding pocket, with contributions from both the SET and SET-I domains (17). Residue Asn101 is positioned near the entrance of this pocket, suggesting a possible role in substrate or cofactor recognition (Figure 2C).

To evaluate the structural consequences of this substitution, an in silico N101S variant was generated based on the available SMYD1 crystal structure (PDB ID: 3n71), followed by local energy minimization (19). In the minimized model, substitution of asparagine with serine introduces a shorter polar side chain with reduced hydrogen bonding capacity and altered geometry. Examination of plausible side chain conformations suggests that this substitution may modify local hydrogen bonding interactions near the entrance of the active site. In particular, the serine hydroxyl group can adopt orientations that favor interactions with nearby secondary structural elements, potentially at the expense of interactions available to the native asparagine residue (Figure 3).

**Figure 3.**
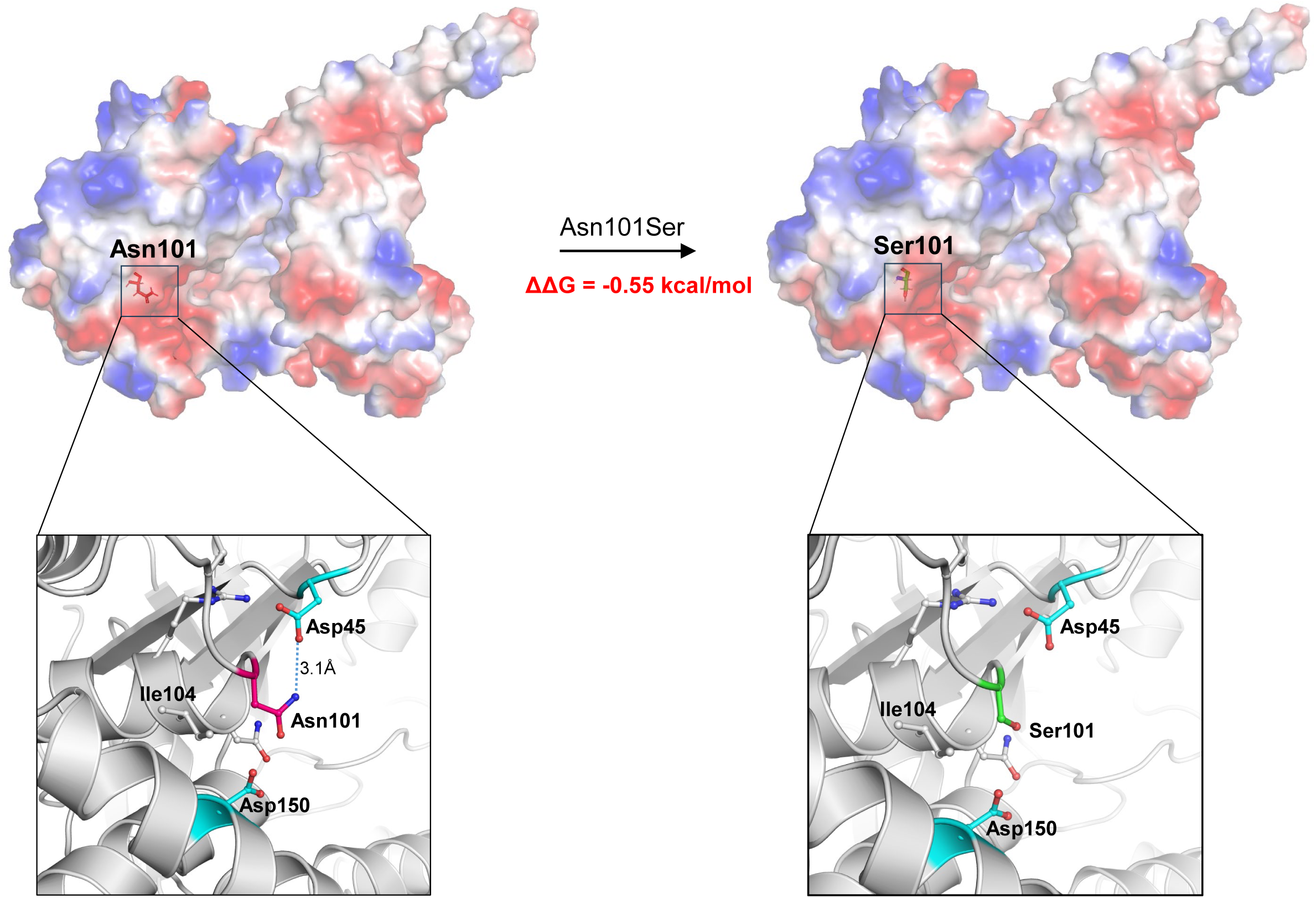
Structural and thermodynamic impact of the N101S substitution in SMYD1. **A.** Electrostatic surface representation of the wild-type SMYD1 structure (PDB ID: 3N71), with Asn101 highlighted (left), and the N101S variant with Ser101 indicated (right). The N101S substitution is predicted to modestly destabilize the protein structure, as indicated by a change in Gibbs free energy (ΔΔG = -0.55 kcal/mol), as calculated using DynaMut2. **B.** Enlarged view of the wild-type structure (left) showing the local interaction network of Asn101, including a hydrogen bond with Asp45 (dashed blue line, 3.1Å), and the N101S variant (right) demonstrating loss of this hydrogen bond with Asp45, resulting in perturbation of the local interaction network near the cofactor binding pocket. All structural models were generated based on the mouse SMYD1 crystal structure (PDB ID: 3N71) and visualized using PyMOL and UCSF Chimera.

To further assess the energetic and dynamic impact of the N101S substitution, we utilized the DynaMut2 web server, which integrates normal mode analysis and graph-based signatures to predict variant-induced changes in protein stability (20). This analysis predicted a stability change (ΔΔG) of -0.55 kcal/mol, indicating a mildly destabilizing effect on the SMYD1 structure. This result supports the notion that the N101S substitution does not induce large-scale structural disruption but may instead lead to subtle local perturbations in stability and flexibility.

Given the proximity of residue 101 to the cofactor binding pocket, such local changes may influence substrate access, positioning, or stabilization within the catalytic site. However, these observations are derived from computational modeling approaches and a locally minimized static structural model, and do not fully account for conformational dynamics or solvent effects. Therefore, while the N101S substitution is predicted to perturb the local interaction network and modestly reduce structural stability, further validation using molecular dynamics simulations or functional assays would be required to establish its impact on SMYD1 enzymatic activity.

### 3.3 *In vitro* evaluation of N101S variant

The highly conserved Asn101 residue is located in the N-terminus of SMYD1 SET-I domain, within close proximity to the substrate binding site. Due to its location and the phenotype of P1, we hypothesized that the N101S variant acts as a hypomorphic allele, resulting in at least partial loss of SMYD1 function. To evaluate the functional significance of this variant, we generated a plasmid containing a missense substitution (c.302A>G) in the mouse ortholog of the human *SMYD1* gene to generate a protein containing the N101S variant. We confirmed the single nucleic acid substitution in the mouse *Smyd1* gene using Sanger sequencing (Figure 4A) as well as the transcript expression of both MYC-tagged *Smyd1*-N101S and wild-type *Smyd1* by RT-qPCR in undifferentiated C2C12 mouse myoblast cells (Figure 4B). The substitution was further validated by immunoblotting, confirming protein expression of both wild-type SMYD1 and MYC-tagged SMYD1-N101S (Figure 4C-D).

**Figure 4.**
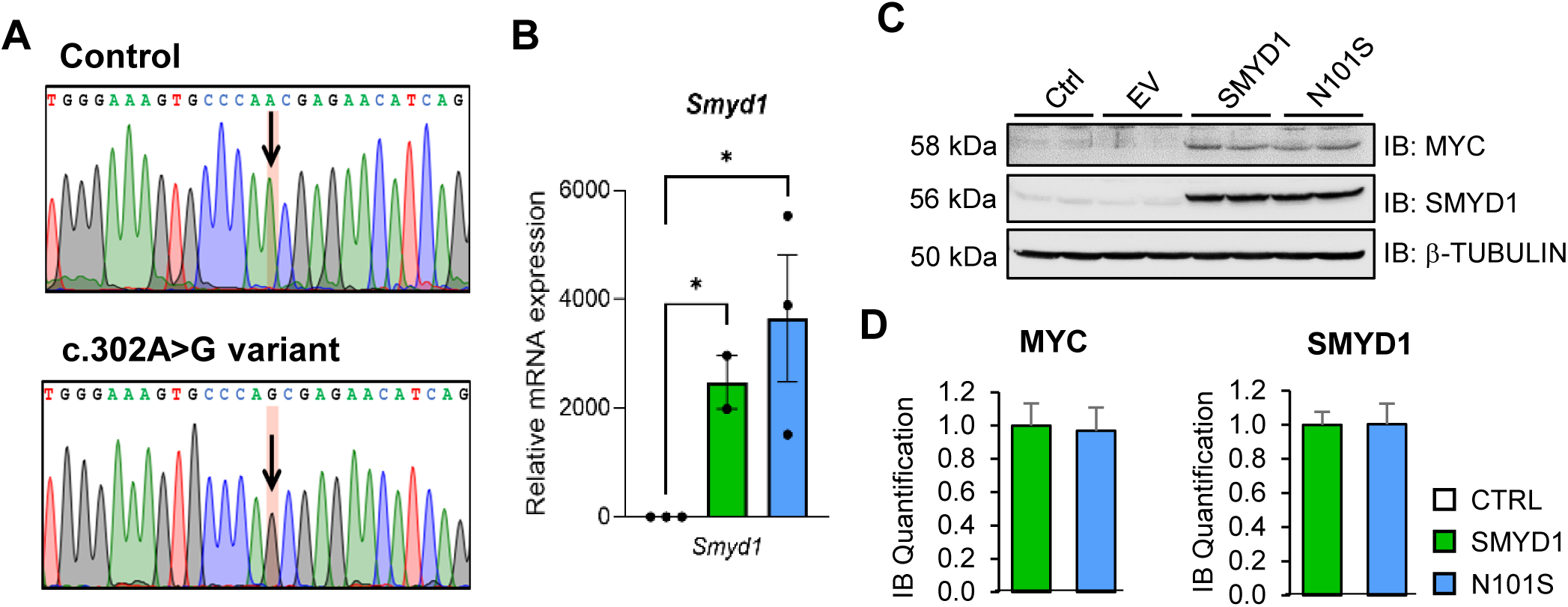
Validation of SMYD1-N101S variant expression in cultured myocytes. Undifferentiated C2C12 cells were transfected with wild-type SMYD1 or N101S constructs or an empty vector (EV) as a control. **A.** Sanger sequencing confirms the construct containing a single nucleotide substitution (A>G) in the mouse *Smyd1* gene at position 302, resulting in a protein carrying the N101S variant. **B.** RT-qPCR analysis confirms increased *Smyd1* transcript expression in both *Smyd1-* and *N101S*-transfected cells. *n*=2-3. Asterisk * indicates *p*<0.05. **C.** Immunoblotting showing expression of both wild-type SMYD1 protein and the MYC-tagged N101S variant, compared to untreated (Ctrl) and empty vector (EV) controls. **D.** Densitometric quantification of immunoblot band intensity for MYC and SMYD1, confirmed in three independent experiments. Data are represented as mean ± SEM.

Through our previous work, we demonstrated that SMYD1 regulates cardiac physiology in part by modulating mitochondrial respiration. To determine whether the N101S variant alters mitochondrial function, we performed a Cell Mito Stress Test using the Seahorse XFe96 Flux Analyzer to measure oxygen consumption rate (OCR) in C2C12 mouse myoblasts expressing either wild-type SMYD1 or N101S variant (Figure 5A). We observed that cells expressing the N101S variant exhibited significantly reduced basal, maximal, and spare respiratory capacity, along with decreased ATP production, compared to wild-type SMYD1 (Figure 5B-C). These findings indicate that the N101S variant impairs mitochondrial respiration *in vitro*.

**Figure 5.**
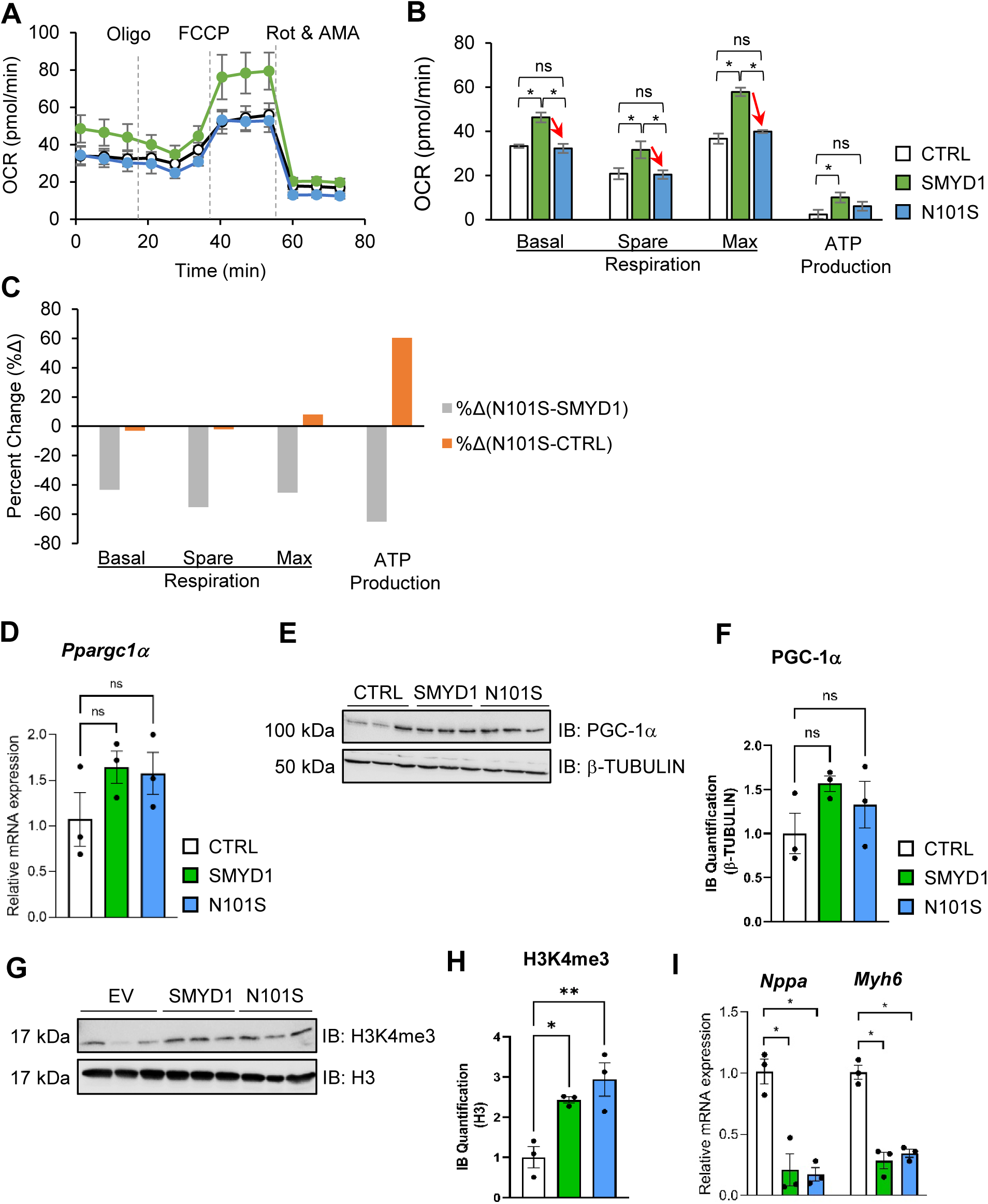
SMYD1-N101S impairs mitochondrial respiratory function in cultured myocytes. C2C12 cells were transfected with wild-type SMYD1 or N101S constructs **or** an empty vector (EV) as a control. **A-C.** Representative traces **(A)**, quantification **(B)**, and percent change **(C)** of mitochondrial oxygen consumption rate (OCR) using a Seahorse Cell Mito Stress Test demonstrating that the N101S variant leads to decreased mitochondrial respiration when compared to wild-type SMYD1 (green). *n*=6. Asterisks: * indicates *p*<0.05, ** indicates *p*<0.01**. D.** RT-qPCR analysis demonstrates no significant change in *Ppargc1α* transcript levels in N101S-expressing cells. *n*=3. **E-F.** Representative immunoblots **(E)** and quantification **(F)** demonstrating no significant change in PGC-1α protein abundance in both wild-type and N101S-expressing cells relative to empty vector control. *n*=3. **G-H.** Representative immunoblots of histone H3K4me3 **(G)** and densitometric quantification **(H)** showing similarly increased H3K4me3 levels in both wild-type and N101S-expressing cells compared to empty vector control, with no significant difference between wild-type SMYD1 and N101S. **I.** RT-qPCR analysis of hypertrophic markers demonstrates no significant change in *Nppa* and *Myh6* transcript levels in N101S-expressing cells relative to wild-type SMYD1. *n*=3. Asterisk * indicates p<0.05. Data are presented as mean ± SEM.

Given that SMYD1 has been shown to regulate mitochondrial energetics in part through its histone methyltransferase-dependent regulation of PGC-1α expression, we next assessed whether this canonical pathway is affected by the N101S variant. In C2C12 cells transfected with wild-type SMYD1 or SMYD1-N101S, we observed no significant differences in *Ppargc1α* transcript levels at 48 h (Figure 5D), nor were there significant changes in PGC-1α protein abundance (Figure 5E-F). Because SMYD1 canonically regulates PGC-1α expression via trimethylation of histone H3K4, we examined whether the N101S variant alters global histone H3K4me3 levels. Immunoblot analysis of total cell lysates revealed that both wild-type SMYD1 and the N101S variant similarly elevated global histone H3K4me3 levels relative to empty vector controls, with no detectable difference between the two constructs. Finally, although C2C12 myoblasts are not a mature cardiomyocyte model, we assessed the expression of the cardiac remodeling-associated genes *Nppa* and *Myh6* as exploratory markers of stress-response transcriptional signaling. Our RT-qPCR analysis showed no significant differences in *Nppa* and *Myh6* transcript levels between cells expressing wild-type SMYD1 and those expressing the N101S variant (Figure 5I), indicating that the mitochondrial defects associated with N101S are not accompanied by detectable changes in these hypertrophy-related transcripts under the conditions tested. Together, these results suggest that the impaired mitochondrial respiration associated with the N101S variant occurs independently of changes in PGC-1α expression or global H3K4me3 levels, indicating an alternative, yet-to-be-identified mechanism underlying the hypomorphic effect of this variant.

### 3.4 Increased cardiac SMYD1 expression in patient with N101S-associated cardiomyopathy

To complement our in vitro findings, we next examined SMYD1 protein expression in cardiac and skeletal muscle tissues from P1, with SMYD1 variant-associated biventricular heart failure at the time of heart transplantation, compared with age- and sex-matched healthy control. Because only a single patient tissue sample was available and no additional patient samples could be obtained for analysis, we performed immunoblotting in triplicate using the same biological sample to assess the technical reproducibility and confirm consistency. Quantification of the three independent immunoblot runs revealed increased SMYD1 protein levels in cardiac tissue from P1 relative to control samples, whereas no difference was observed in skeletal muscle (Figure 6A-B). The data suggest a cardiac tissue-specific upregulation of SMYD1 protein levels in the failing heart of P1, which may represent a compensatory response to the reduced SMYD1 activity conferred by the N101S variant. Despite increased protein abundance, the presence of the N101S variant may impair SMYD1-dependent regulation of downstream substrates and pathways. Further studies will be needed to define how the N101S variant alters SMYD1 substrate specificity and downstream epigenetic regulation in cardiac tissue.

**Figure 6.**
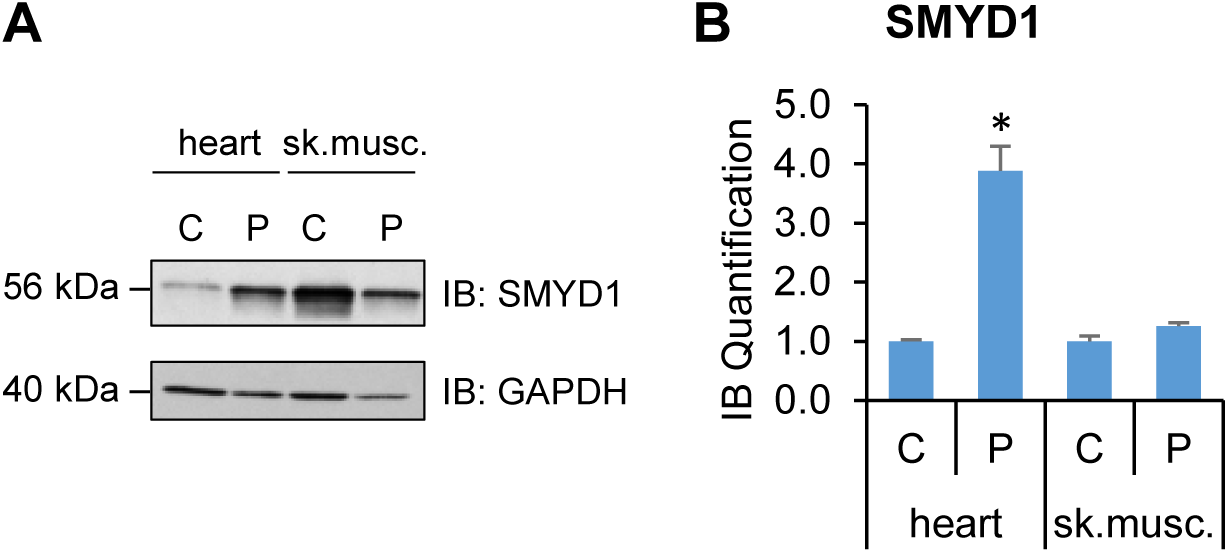
Increased SMYD1 protein abundance in cardiac but not skeletal muscle tissue from a patient with SMYD1 variant-associated biventricular heart failure. **A.** Representative immunoblots showing SMYD1 protein levels in cardiac and skeletal muscle tissues obtained from patient P1 at the time of heart transplantation, compared with an age- and sex-matched healthy control. GAPDH was used as a loading control. **B.** Densitometric quantification of SMYD1 protein abundance normalized to GAPDH, shown as relative protein expression. SMYD1 protein abundance was increased in cardiac tissue from P1 relative to the healthy control, whereas no significant difference was observed in skeletal muscle. C: healthy donor control; P: patient P1; sk.musc.: skeletal muscle. Asterisk * indicates *p*<0.05. Data are presented as mean ± SEM.

## 4 Discussion

Infantile cardiomyopathy represents a genetically heterogeneous group of disorders in which the identification of causal variants has been greatly accelerated by next-generation sequencing approaches such as WES (24). However, translating genetic findings into a mechanistic understanding remains a critical gap, particularly for rare or novel variants in genes with limited human data.

In this study, we investigated the functional consequences of the SMYD1 c.302A>G (p.Asn101Ser) variant identified in a previously reported patient with severe early-onset cardiomyopathy. While our prior reports established the clinical phenotype and genetic findings, the impact of this variant at the molecular level had not been explored. We provide evidence supporting a hypomorphic effect of the N101S substitution through combined structural, *in vitro,* and human tissue analyses.

Our molecular modeling analysis predicts that the N101S substitution introduces subtle but functionally relevant perturbations in the local structural environment of SMYD1. The Asn101 residue is highly conserved and positioned near the cofactor-binding pocket within the SET-I domain. Substitution to serine results in reduced side chain length and loss of a hydrogen bond with the neighboring Asp45 residue, which is predicted to modestly destabilize the protein (ΔΔG = -0.55 kcal/mol). While this predicted destabilization is modest, even small structural perturbations in catalytically important regions can significantly alter enzymatic activity. Given the high sequence homology between mouse and human SMYD1, these structural insights are likely to be biologically relevant.

Consistent with these predictions, our in vitro studies demonstrate that the N101S variant impairs mitochondrial respiratory capacity. Cells expressing the N101S variant exhibit reduced basal and maximal respiration, as well as decreased ATP production, indicating compromised mitochondrial bioenergetics. These findings are consistent with prior work demonstrating that SMYD1 is a key regulator of mitochondrial function in cardiomyocytes (9, 21). Notably, this impairment occurred independently of changes in PGC-1α expression or global H3K4me3 levels, suggesting that the N101S variant disrupts mitochondrial function through a non-canonical, PGC-1α-independent mechanism. This is a novel finding that distinguishes the hypomorphic effect of the N101S variant from the complete loss of SMYD1 function observed in knockout models, where PGC-1α downregulation is a primary downstream consequence (9). The identity of the relevant non-histone substrate or alternative pathway through which SMYD1 regulates mitochondrial function in the context of the N101S variant remains an important open question for future investigation.

Interestingly, analysis of cardiac tissue from the proband revealed increased SMYD1 protein abundance, compared to healthy controls, without corresponding changes in skeletal muscle. This suggests a tissue-specific upregulation of SMYD1 in the failing heart. Given the functional impairment observed *in vitro*, this increase is most consistent with a compensatory response to reduced SMYD1 activity rather than restoration of downstream signaling. However, whether elevated SMYD1 levels preserve its epigenetic and mitochondrial regulatory functions in the presence of the N101S variant remains unclear and warrants further investigation.

Although SMYD1 has been extensively studied in model organisms, reports of human disease-associated variants remain limited. The first reported variant (c.814T>C; p.Phe272Leu), identified in a patient with hypertrophic cardiomyopathy, was predicted to be deleterious by in silico tools but lacked comprehensive functional validation (13). A second variant (c.675delA; p.Lys225Asnfs*8), identified in a patient with left ventricular non-compaction cardiomyopathy, is predicted to result in loss of function due to truncation of the protein, although direct experimental validation has not been reported (14). It is worth noting that both previously reported variants are heterozygous with limited functional studies, whereas the N101S variant in P1 is homozygous. This study expands this limited body of evidence by providing functional characterization of a third SMYD1 variant and directly linking impaired mitochondrial respiration to SMYD1 dysfunction in human cardiomyopathy.

Consistent with the clinical and histopathological findings previously reported in P1, including the structural abnormalities in myofibrillar organization and mitochondrial architecture (16), our current findings further support a role for SMYD1 in both cardiac and skeletal muscle biology. These pathological features are also consistent with prior studies in zebrafish and mouse models, where SMYD1 has been shown to regulate sarcomere assembly and mitochondrial function (25–27). Interestingly, despite these structural abnormalities, the proband did not exhibit significant skeletal muscle weakness during early childhood. This may suggest differential functional requirements for SMYD1 in cardiac versus skeletal muscle, or alternatively, the presence of compensatory mechanisms in skeletal muscle that are absent in the heart. Whether skeletal muscle dysfunction manifests later in life in patients with SMYD1 variants remains an open clinical question.

The observed mitochondrial abnormalities (16) are particularly notable in the context of early postnatal cardiac development, a period characterized by a metabolic shift toward oxidative phosphorylation and fatty acid β-oxidation (28, 29). Disruption of mitochondrial function during this critical window may contribute significantly to disease severity. In this context, the impaired mitochondrial respiration observed in cells expressing the N101S variant provides a plausible mechanistic link between the molecular defect and the clinical phenotype.

Several limitations of this study deserve acknowledgment. First, functional studies were performed in undifferentiated C2C12 mouse myoblasts that do not express SMYD1 endogenously, rather than cardiomyocytes, and further validation in iPSC-derived cardiomyocytes or neonatal rat ventricular cardiomyocytes will be important to confirm these findings in a cardiac-specific context. Second, this study is based on a single patient, which is inherent to the rarity of this condition but limits generalizability. Third, P1 was found to also carry a homozygous pathogenic *MYBPC3* variant (c.1224-52G>A), which is independently known to cause severe cardiomyopathy. While the functional data presented here support a direct role for the SMYD1 N101S variant in mitochondrial impairment, the relative contribution of each variant to the overall clinical phenotype cannot be fully resolved from the current data.

In summary, our results support that the SMYD1 N101S variant is a destabilizing hypomorphic variant that impairs mitochondrial function through a novel PGC-1α-independent mechanism (Figure 7). This study provides mechanistic insight into SMYD1-associated cardiomyopathy and contributes to the growing understanding of how genetic variation in this gene affects cardiac biology. The identification of the downstream alternative pathways through which SMYD1 regulates mitochondrial bioenergetics represents an important direction for future work, with potential implications for understanding and treating SMYD1-associated cardiomyopathies.

**Figure 7.**
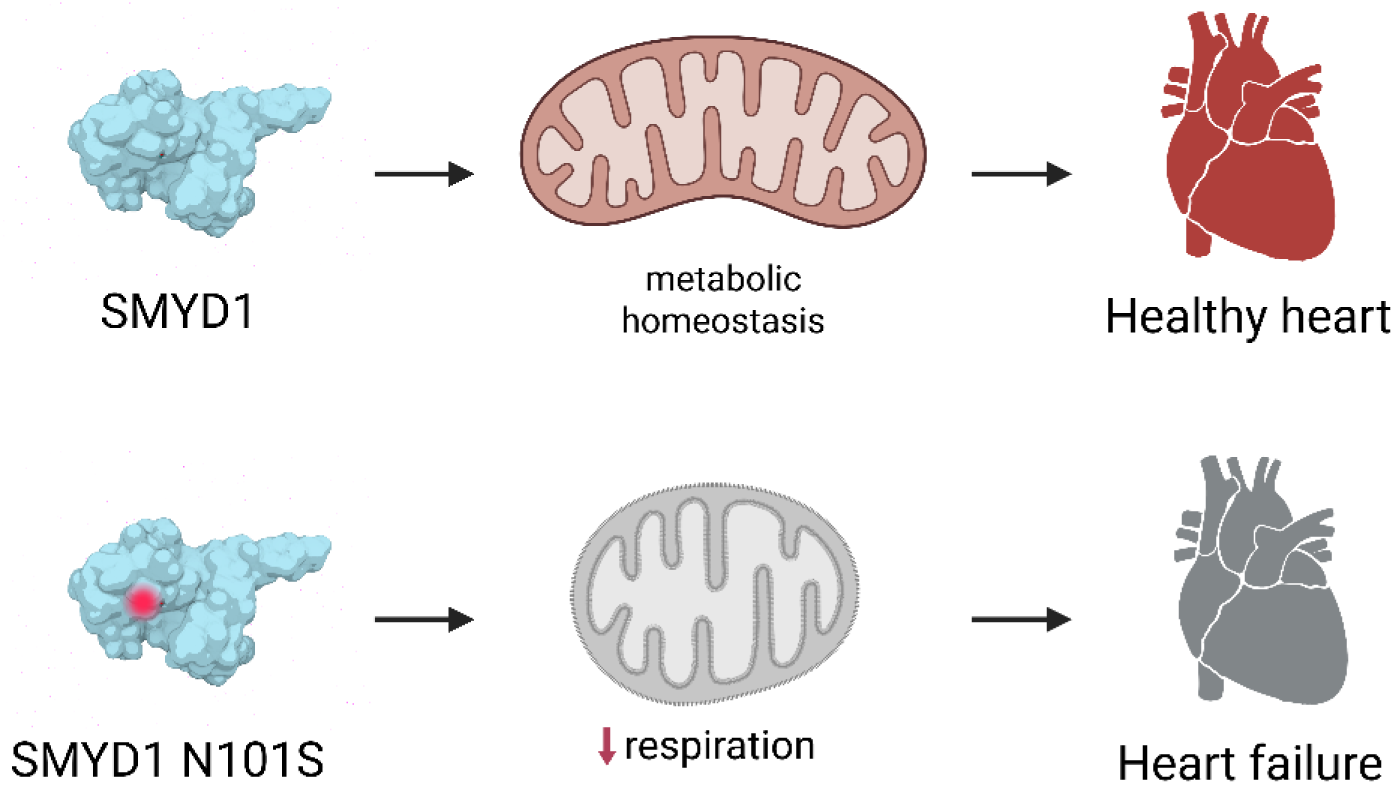
Proposed mechanism of SMYD1 N101S variant dysfunction in the heart. Upper panel: Wild-type SMYD1 supports mitochondrial function and maintains metabolic homeostasis in the heart, contributing to normal cardiac function. **Lower panel:** the N101S variant impairs mitochondrial respiratory capacity, leads to compromised metabolic homeostasis and heart failure. The red circle on the SMYD1 N101S protein indicates the location of the N101S substitution within the SET-I domain.

## Acknowledgments

The authors are grateful to the family who agreed to participate in this research study.

We thank Dr. Ying Li and the Metabolic Phenotypic Core at the University of Utah for assistance in Seahorse OCR measurements. This work was supported by the Nora Eccles Harrison Treadwell Foundation Grant 10038331 and the National Institutes of Health Grants R01-HL-130424 and F32-HL-144034. The contents of this manuscript are solely the responsibility of the authors and do not necessarily represent the official view of the NIH.

## Author contributions

MWS and SF concept and design of manuscript; MWS and CG performed experiments in cell culture; MC helped with site-directed mutagenesis, MWS performed structural modeling; LGG performed clinical evaluation of patients and provided clinical description; LGG obtained consent from the patient’s parents, collected samples, and completed genetic testing on the rest of the family members; MWS, prepared figures; MWS drafted the manuscript; MWS, CG, SF and LGG edited and revised the manuscript and figures; MWS, SF, and LGG analyzed data and interpreted results. All authors read and approved the final version of the manuscript.

